# Basal-to-classical phenotypic switch of pancreatic cancer cells upon integration into the duodenal epithelium

**DOI:** 10.1101/2021.02.01.429212

**Authors:** Benedek Bozóky, Carlos Fernández Moro, Carina Strell, Ingemar Ernberg, Laszlo Szekely, Marco Gerling, Béla Bozóky

**Affiliations:** Theme Cancer, oncology, Karolinska University Hospital, 17 176 Solna, Sweden; Department of Microbiology, Tumor, and Cell Biology, Karolinska Institutet, Solnavägen 9, 171 65 Solna, Sweden; Department of Clinical Pathology/Cytology, Karolinska University Hospital, Stockholm, 14186, Sweden; Department of Laboratory Medicine, Division of Pathology, Karolinska Institutet, 14186 Stockholm, Sweden; Department of Oncology-Pathology, Karolinska Institutet, 17164 Solna, Sweden; Department of Immunology, Genetics and Pathology, Uppsala University, 75185 Uppsala, Sweden; Department of Biosciences and Nutrition, Karolinska Institutet, 14183 Huddinge, Sweden

## Abstract

Pancreatic ductal adenocarcinoma (PDAC) is one of the most aggressive solid tumors. Based on transcriptomic classifiers, basal-like and classical PDAC subtypes can be defined that differ in prognosis. Single-cell sequencing has recently revealed that these subtypes coexist in individual tumors. However, the contribution of either clonal heterogeneity or microenvironmental cues to subtype heterogeneity is unclear. Here, we report the tumor phenotype dynamics in a cohort of patients in whom PDAC infiltrated the duodenal wall. Using multiplex immunohistochemistry, we show that PDAC cells revert to non-destructive growth and undergo differentiation towards the classical subtype upon integration into the duodenal epithelium. Our results tightly link microenvironmental cues to the PDAC molecular subtype and open the door to a systematic investigation of microenvironmental control in human pancreatic cancer.

## Introduction

Pancreatic ductal adenocarcinoma (PDAC) is one of the most lethal tumors with a five-year survival rate of less than 9% (1). Although clinically perceived as uniformly aggressive, PDAC subtypes can be defined based on transcriptional profiling (2–6). Most classifications, including the most recent one, distinguish classical from basal-like tumor subtypes, although other subtypes and intermediate states exist (2). Basal-like tumors are uniformly associated with an inferior prognosis (2–4, 6). While subtyping based on bulk transcriptomic data is prognostically valuable, both basal-like and classical cells exist in individual tumors, as revealed by single-cell RNA sequencing (2). Genetic changes, such as allelic imbalances of mutant *KRAS*, contribute to this intratumor heterogeneity (2, 7); however, mouse models have demonstrated a central role for the microenvironment in shaping the PDAC tumor cell phenotype (8), and there is a strong support from experimental studies that the non-malignant microenvironment can reprogram malignant cells to a normal-like behavior, associated with tumor cell differentiation (9, 10). In human PDAC, cancer cell states have recently been linked to adjacent fibroblast subtypes (11); nevertheless, the full extent to which microenvironmental cues shape tumor cell phenotypes remains unclear, and the forces driving tumor cell heterogeneity have not been well defined.

Morphological heterogeneity of pancreatic tumors is regularly observed during routine pathological assessment, which is the main pillar of PDAC diagnosis (12). Based on histopathology, it has previously been reported that PDAC cancer cells can appear more differentiated upon entering the duodenum, which occasionally leads to misdiagnosis of PDAC as an intestinal neoplasm (13). From a tumor biology perspective, these striking phenotypical changes in tumor cells upon switching location provide a remarkable, yet largely overlooked, possibility of studying cancer cell plasticity in relation to microenvironmental cues within spatially defined tissue compartments in humans.

Here, we set out to systematically map the tumor cell phenotype dynamics in a unique cohort of PDAC patients with duodenal tumor infiltration. We find that PDAC cells revert to non-destructive growth and undergo classical/intestinal differentiation upon integration into the duodenal epithelium. Our results link the intestinal microenvironment to defined shifts in the PDAC cancer cell phenotype.

## Results

We studied n = 20 patients with infiltration of PDAC cells through the duodenal wall and into the intestinal epithelium. We observed consistent morphological changes in areas of PDAC integration into the epithelial layer (**Figure 1A**). The mucosal architecture was strikingly preserved, and the epithelial lining exhibited either normal morphology or mild reactive atypia, interspersed with areas of columnar epithelium with a dysplastic/neoplastic appearance. To confirm unequivocally that the neoplastic cells in the duodenal mucosa were of pancreatic origin, rather than reactive intestinal cells secondary to PDAC infiltration into the submucosa, we used two independent immunohistochemical markers, SMAD4 and p53; SMAD4 is lost in approximately half of all pancreatic cancer cases (14); while *TP53* mutations, which lead to accumulation of mutant p53 protein, are detectable in 50% of patients, independently of alterations in *SMAD4* (14). Both markers confirmed the seamless integration of PDAC cells into the SMAD4^+ve^/p53^−ve/low^ duodenal epithelium, without destruction of the mucosal architecture (**Figures 1B&C**). Intramucosal PDAC cells were well differentiated and polarized, in contrast to the submucosa, which harbored pleomorphic cancer cells and irregular glands, consistent with primary PDAC histomorphology (**Figures 1D&E, same tumors as in B&C**).

**Figure 1:**
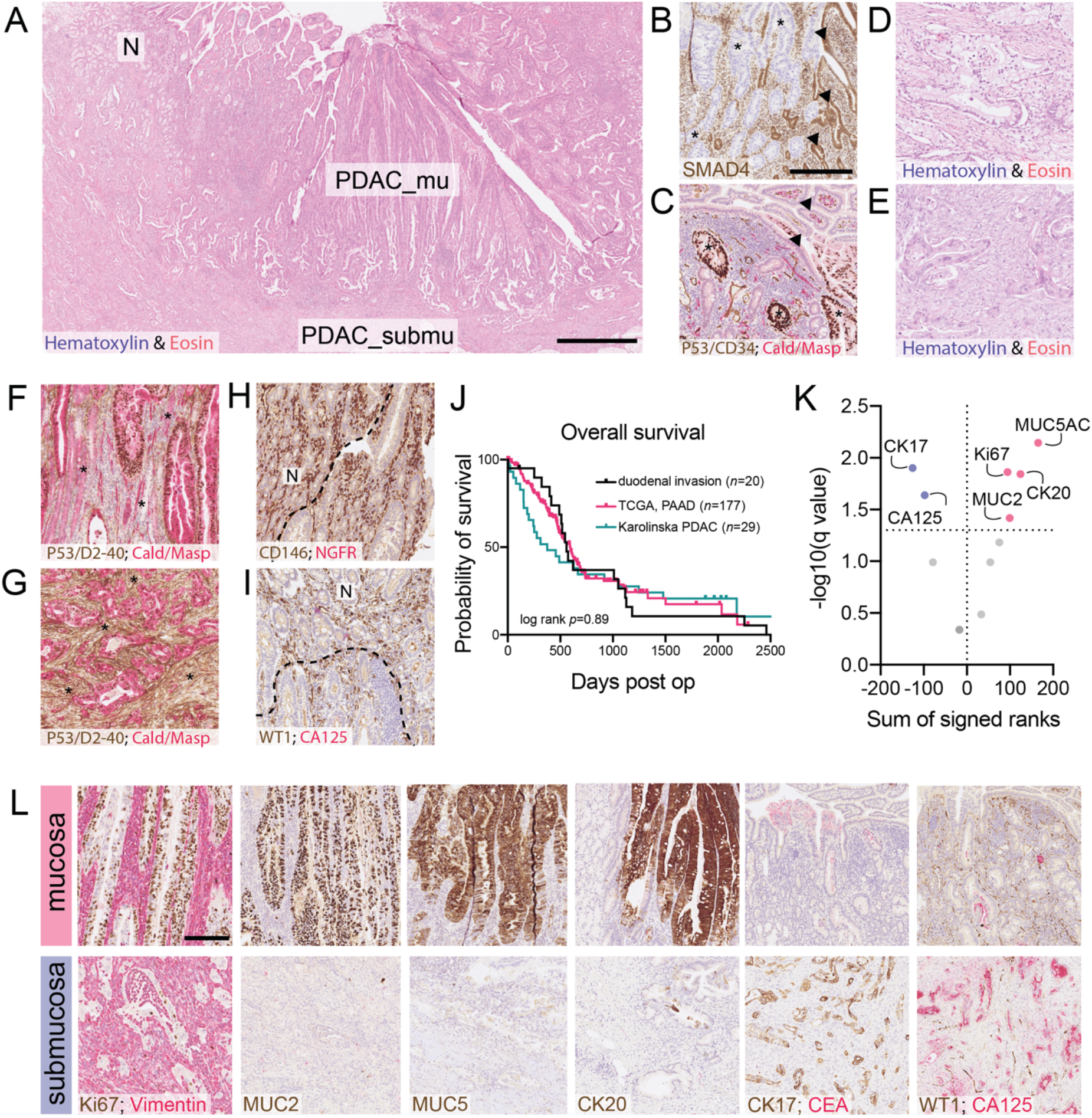
Pancreatic ductal adenocarcinoma infiltrating the duodenal mucosa. **A)** Hematoxylin & Eosin (H&E) staining of pancreatic ductal adenocarcinoma (PDAC) cells that have invaded and integrated into the duodenal mucosa. N: non-neoplastic duodenal epithelium, PDAC_mu: region of PDAC cells within the mucosa, PDAC_submu: region of PDAC cells in the submucosa. Scale bar: 1 mm. **B)** Representative immunohistochemistry (IHC) for SMAD4 in a PDAC with genetic loss of *SMAD4* shows intestinal villi lined by SMAD4-negative PDAC cells (asterisks) compared to adjacent small intestinal epithelial cells positive for SMAD4 expression (arrowheads). Scale bar: 250 μm, applies to (B)–(I). All IHC counterstained with Hematoxylin. Multiplex staining combinations are indicated in panels, text color denotes color of chromogen used to visualize protein expression. **C)** Representative image of p53 IHC in a PDAC with accumulation of p53 protein due to *TP53* mutation; note regions of p53-positive cells (PDAC cells, asterisks) adjacent to p53-negative cells (intestinal epithelial cells, arrowheads). **D)** and **E)** H&E staining of PDAC cells in the submucosa: (D) is the same tumor as in (B), (E) is the same tumor as in (C). **F)** and **G)** Quadruple IHC for the indicated proteins illustrating D2-40 overexpression as a marker for desmoplasia in the submucosa (G), but absent in the mucosa (F); note that in these quadruple stainings, brown marks both D2-40 (stromal) and p53 (epithelial), and red marks both caldesmon (stromal) and maspin (epithelial); asterisks indicate stroma. **H)** IHC for the *lamina propria* marker, CD146, shows preserved expression in areas of intraepithelial tumor integration. **I)** IHC for another *lamina propria* marker, WT1, also shows preserved expression; dotted lines in (H) and (I) represent approximate border between PDAC cells and enterocytes; (B)–(I) present representative staining of *n* ≥ 10 cases. **J)** Comparison of overall survival in the current cohort of patients with duodenal infiltration *vs*. patients included in a previously published cohort operated on for PDAC at the same center (15), and *vs*. patients from The Cancer Genome Atlas (TCGA) PAAD dataset. Initial patient numbers and log-rank *p* given in panel. **K)** Volcano plot showing significant findings in red (higher in mucosa) and blue (higher in submucosa out of a total of n = 12 markers included in analysis as indicated in the text. Data based on results from Wilcoxon matched-pairs signed rank test, multiple test correction with two-stage step-up (ref. 16), fdr < 0.05. **L)** Representative images of the indicated markers showing significant differential expression in mucosal *vs*. submucosal tumor cells. Scale bar: 250 μm, applies to all images in (L); representative staining of n ≥ 15 cases. Note that multiplex IHC was performed for some markers (e.g. Ki67 together with vimentin in the left panel), but not all markers were included in quantitative analysis (see *Methods* for details).

A hallmark of PDAC is its desmoplastic stroma, in which tumor glands are embedded (11). Notably, IHC for the desmoplasia marker, podoplanin (D2-40), revealed no desmoplasia of the subepithelial stroma adjacent to mucosal cancer cell integration (**Figure 1F**), while desmoplasia was present in the submucosa (**Figure G**). In contrast, expression of CD146 and WT1, markers of the intestinal *lamina propria* (see also *Methods*), was preserved in regions adjacent to intraepithelial tumor cells (**Figures 1H&I**).

There was no difference in overall survival (OS) in our group of 20 patients compared to PDAC patients operated on at the same hospital (15), or to PDAC cases from the TCGA cohort (**Figure 1J**). Hence, the data indicate that duodenal mucosal infiltration does not define a prognostically distinct patient subgroup.

The diverging phenotypes of PDAC cells in mucosal *vs*. submucosal locations implies a high degree of location-dependency, suggesting that the local microenvironment is a major contributor to the tumor cell phenotype. To quantify these phenotypic changes, we used serial multiplex quantitative immunohistochemistry (smq-IHC) to assess a panel of 12 protein markers, selected to identify tumor cell characteristics, and to approximate the transcriptional subtypes (see also *Methods*) (2). The panel comprised intestinal and pancreatobiliary differentiation markers (CK20, CDX2, MUC2; MUC1, MUC5AC), all of which are included in the classical transcriptional profile (2), a pancreatobiliary marker specific to the basal-like profile (CK17), general markers for PDAC cells (CK7, MUC6) and tumor-specific glycoproteins (CA19-9, CA125, and CEA) together with Ki67 to assess proliferation.

In agreement with the phenotypic shift, smq-IHC revealed significantly reduced expression of the basal/pancreatobiliary markers, CK17 and CA125, in the mucosa *vs.* submucosa, while expression of the classical/intestinal markers, MUC5AC, CK20, and MUC2 was increased (**Figures 1K&L**), supporting a strong switch from basal-like towards classical/intestinal differentiation upon intramucosal integration, accompanied by enhanced Ki67 positivity (**Figures 1K&L**).

## Discussion

The integration of PDAC cells into the duodenal mucosa leads to a quantifiable phenotypic shift towards intestinal differentiation. Location-dependent morphological changes are accompanied by a loss of basal-like markers in favor of classical markers, corresponding to a switch towards a less aggressive molecular phenotype (2–4, 6). Interestingly, mucosal PDAC cells were cycling (Ki67^+ve^), suggesting the uncoupling of differentiation from proliferation. Together, the data define a real-life endpoint of the phenotypic plasticity of PDAC cells in humans. Although conjectural, they strongly support a model in which basal-like *vs*. classical tumor subtypes are tightly influenced by microenvironmental cues. The consistency of this phenomenon suggests that this and similar cohorts displaying duodenal invasion can be invaluable for deciphering the molecular underpinnings of PDAC subtype emergence orchestrated by the microenvironment.

## Material and Methods

### Patients

Patients were identified by a retrospective search in the pathology archive database of the Karolinska University Hospital, Huddinge, Sweden, and through routine diagnostic pathology. Cases in which infiltration of PDAC cancer cells into the small intestine was described in the pathology report or for which the participating pathologists (CFM, Béla B, LS) noted this phenomenon were selected and systematically reassessed based on available Hematoxylin and Eosin staining and routine immunohistochemistry (IHC). Patients operated on between 2008 and 2020 were considered. Clinical data were obtained by retrospective chart review. A total of n = 20 patients in whom PDAC cells had infiltrated the entire thickness of the duodenal wall were included. All patients (n = 8 females, n = 12 males, aged 64–82 years, median age 71.8 years) underwent resection according to Whipple at the Karolinska University Hospital, Stockholm, Sweden between 2008 and 2020.

### Other Cohorts

A local cohort of histologically confirmed PDAC cases that underwent Whipple’s procedure between 2008 and 2016 has been described previously (15). This cohort was curated to remove patients overlapping with the current cohort, leaving n = 29 patients for comparison. The Cancer Genome Atlas (17) PAAD cohort was accessed via https://portal.gdc.cancer.gov/ and curated to remove neuroendocrine tumors, leaving n = 177 cases for survival analysis. Statistical analyses were performed with GraphPad prism v9.0.0. The study was approved by the responsible ethical review boards.

### Serial multiplex quantitative immunohistochemistry

smq-IHC (**Figure 2)** was performed as described previously (15). Antibodies are detailed in **Table 1**. Formalin-fixed paraffin-embedded (FFPE) samples were cut to a thickness of 4 μm and stained on an automated stainer (BOND-MAX, Leica Biosystems, Germany) as part of the diagnostic routine in a clinically accredited histology lab. Staining procedures have been described previously (15). All antibodies (**Table 1**) were extensively validated for clinical use. For all antibody stainings, at least n = 10 cases were included for the final assessment based on staining quality and availability. For all quantified stainings (**Figure 1K and L**), statistics are based on the evaluation of at least n = 15 matched mucosa/submucosa pairs.

**Figure 2:**
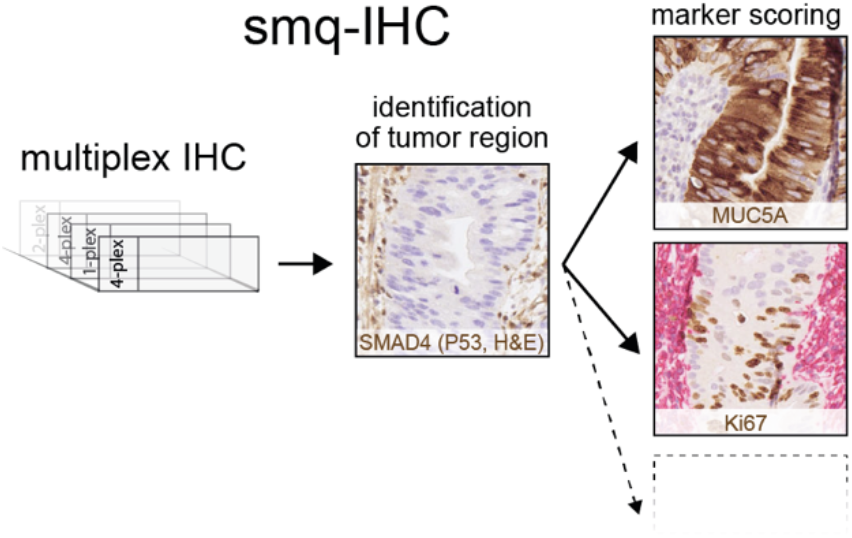
Schematic of serial multiplex quantitative immunohistochemistry (smq-IHC). Serial sections are stained with different antibody combinations. IHC of SMAD4 and p53 guides identification of tumor regions. Carefully matched regions of serial sections are evaluated, while SMAD4/p53 sections are continuously used to navigate.

---

**Supplementary Table 1.** Antibodies used in this study. Pretreatment “H1”: Bond Epitope Retrieval Solution 1, citrate, 20 minutes,
“H2” = Bond Epitope Retrieval Solution 2, EDTA, 20 minutes; NCL: Novocastra/Leica.

| *antigen* | *full name* | *main target cell* | *clone* | *dilution* | *pretreatment* | *manufacturer, cat nr* |
| --- | --- | --- | --- | --- | --- | --- |
| Caldesmon | Caldesmon | Smooth muscle cells (18) | H-CD | 1:300 | H2 | Dako-M3557 |
| CD146 | Cluster of differentiation 146 | *Lamina propria* stroma, pericryptal mesenchymal cells (19) | UMAB154 | 1:100 | H2 | Origene-UM800051 |
| CD34 | Cluster of differentiation 34 | Fibrocytes/non-myofibroblastic stromal cells of the *lamina propria* (20, 21) | QBEnd/10 | 1:50 | H2 | Dako-M7165 |
| CK7 | Cytokeratin 7 | PDAC cells (pan PDAC marker) (22) | RN7 | 1:200 | H2 | NCL-L-CK7-560 |
| CK17 | Cytokeratin 17 | PDAC cells (15), basal-like type (2, 3) | E3 | 1:25 | H2 | NCL-CK17 |
| CK20 | Cytokeratin 20 | cells with intestinal differentiation (23), classical type (2, 3) | PW31 | 1:25 | H2 | NCL-L-CK20-561 |
| D2-40 | Podoplanin | desmoplastic stroma (24) | D2-40 | 1:50 | H2 | Dako-M3619 |
| vimentin | Vimentin | stromal cells, cells undergoing EMT (25, 26) | V9 | 1:1500 | H1 | Dako-M0725 |
| MUC1 | Mucin 1 | PDAC cells (27) | Ma695 | 1:50 | H1 | NCL-MUC1 |
| MUC2 | Mucin 2 | intestinal goblet cells (28), PDAC cells of classical type (2) | Ccp58 | 1:100 | H2 | NCL-MUC2 |
| MUC5AC | Mucin 5AC | PDAC cells (15), classical type (2) | CLH2 | 1:50 | H2 | NCL-MUC-5Ac |
| MUC6 | Mucin 6 | PDAC cells, pancreatic differentiation (15) | CLH5 | 1:50 | H2 | NCL-MUC-6 |
| NGFR | Nerve growth factor receptor (p75) | Pancreatic stroma (29) | "Polyconal" | 1:200 | H2 | Atlas-HPA004765 |
| CEA(-m) | Carcinoembryonic antigen - monoclonal | PDAC cells (15) | 11-7 | 1:400 | H2 | Dako-M7072 |
| CA125 | Cancer-antigen 125 | PDAC cells (15) | OV185:1 | 1:100 | H2 | NCL-L-Ca125 |
| CA19-9 | Cancer-antigen 19-9 | PDAC cells (30) | CA241:5:1:4 | 1:400 | H2 | NCL-L-CA19-9 |
| Maspin | Maspin | PDAC cells (31) | EAW24 | 1:50 | H2 | NCL-Maspin |
| WT1 | Wilms tumor 1 (WT1 transcription factor) | cells of the intestinal *lamina propria* (32) | 6F-H2 | 1:100 | H2 | Dako-M3561 |
| CDX2 | caudal type homeobox 2 | intestinal epithelial cells (33), PDAC cells of classical type (2) | AMT28 | 1:25 | H2 | NCL-CDX2 |
| p53 | tumor protein p53 | p53 protein (34) | Do-7 | 1:300 | H1 | NCL-L-P53-D07 |
| Ki67 | marker of proliferation Ki-67 | proliferating cells (35) | MIB-1 | 1:150 | H2 | Dako-M7240 |
| SMAD4 | Mothers against decapentaplegic homolog 4 | SMAD4 protein (36) | B-8 | 1:300 | H2 | Santa Cruz-sc-7966 |
| Chromogranin A | Chromogranin A | enteroendocrine cells (37) | 5H7 | 1:100 | H1 | NCL-CHROM-430 |

## Acknowledgements

Marco Gerling’s research group is supported by The Swedish Research Council, The Swedish Society for Medical Research, the Åke Wiberg Foundation, the Jeansson Foundation and the Karolinska Institute. Ingemar Ernberg’s laboratory is supported by The Swedish Cancer Society. We are thankful for the support provided by the histological and immunohistochemical laboratories, Karolinska University Hospital, and to Rune Toftgård for comments on the manuscript. The results shown here are partly based upon data generated by the TCGA Research Network: https://www.cancer.gov/tcga.

